# Intestine-specific deletion of metal transporter *Zip14 (Slc39a14)* causes brain manganese overload and locomotor defects of manganism

**DOI:** 10.1101/2020.01.10.902122

**Authors:** Tolunay B. Aydemir, Trista L. Thorn, Courtney H. Ruggiero, Marjory Pompilus, Marcelo Febo, Robert J. Cousins

**Author notes:** **Corresponding author:** Tolunay Beker Aydemir, Ph.D., Cornell University, Division of Nutritional Sciences, Ithaca, NY, 14853.

## Abstract

Impaired manganese (Mn) homeostasis can result in excess Mn accumulation in specific brain regions and neuropathology. Maintaining Mn homeostasis and detoxification is dependent on effective Mn elimination. Specific metal transporters control Mn homeostasis. Human carriers of mutations in the metal transporter ZIP14 and whole-body *Zip14* KO (WB-KO) mice display similar phenotypes, including spontaneous systemic and brain Mn overload, and motor dysfunction. Initially, it was believed that Mn accumulation due to *ZIP14* mutations caused by impaired hepatobiliary Mn elimination. However, liver-specific *Zip14* KO mice (L-KO) did not show systemic Mn accumulation or motor deficits. ZIP14 is highly expressed in the small intestine and is localized to the basolateral surface of enterocytes. Thus we hypothesized that basolaterally-localized ZIP14 in enterocytes provides another route for elimination of Mn. Using wild type and intestine-specific ZIP14 KO (I-KO) mice, we have shown that ablation of intestinal *Zip14* is sufficient to cause systemic and brain Mn accumulation. The lack of intestinal ZIP14- mediated Mn excretion was compensated for by the hepatobiliary system; however, it was not sufficient to maintain Mn homeostasis. When supplemented with extra dietary Mn, I-KO mice displayed some motor dysfunctions, brain Mn accumulation based on both MRI imaging and chemical analysis, thus demonstrating the importance of intestinal ZIP14 as a route of Mn excretion. A defect in intestinal *Zip14* expresssion likely could contribute to the Parkinson-like Mn accumulation of manganism.

**New & Noteworthy:** Mn-induced parkinsonism is recognized as rising in frequency due to both environmental factors and genetic vulnerability, yet currently, there is no cure. We provide evidence in an integrative animal model that basolaterally localized ZIP14 regulates Mn excretion and detoxification and that deletion of intestinal ZIP14 leads to systemic and brain Mn accumulation, providing robust evidence for the indispensable role of intestinal ZIP14 on Mn excretion.

## Introduction

Accumulation of manganese (Mn) in the brain of rodents produces a neurodegenerative disease (manganism) with signatures that are similar to those in humans with Parkinson’s disease-like symptoms (parkinsonism) and dystonia (26, 27, 30, 32). Recently, considerable attention has been given to Mn-induced neurodegeneration and parkinsonism in humans (1, 10, 15, 22, 24). Potential causes of excess Mn accumulation include occupational exposure and contaminated drinking water (36), vegetarian diets and overconsumption of Mn-containing dietary supplements (12), chronic liver disease (28, 34), and abuse of amphetamine-like drugs (13). The deleterious effects of Mn exposure on human health are amplified by the increased atmospheric levels of Mn from exhaust emissions (8). These numerous potential routes of chronic Mn exposure, coupled with increased lifespans of humans, have increased the prevalence of Mn-related neurodegenerative disease (ND) worldwide (9, 11, 18, 29).

Mutations in the human metal transporter genes *ZNT10*, *ZIP8*, and *ZIP14* impair Mn homeostasis (6, 19, 20, 25, 28, 35). Mutations in the *SLC39A14* gene encoding the ZIP14 transporter were found in patients with childhood-onset parkinsonism and dystonia with systemic and brain Mn accumulation (35). Since that discovery, there have been a number of case reports of similar symptoms in humans with *ZIP14* mutations (17, 23, 31, 38). Whole-body *Zip14* KO mice display phenotypes similar to those of human carriers of *ZIP14* mutations, including spontaneous systemic and brain Mn overload, specifically in the globus pallidus, and motor dysfunction (3). Hepatic Mn uptake and intestinal Mn elimination were also impaired in whole-body *Zip14* KO mice compared with WT mice.

Mn elimination is crucial for maintaining homeostasis and detoxification; however, very little is known about how it is regulated. ZIP14 is highly expressed in the small intestine and liver at steady-state compared with other tissues (21). The main route of Mn excretion is via bile (15). Thus, Mn accumulation due to ZIP14 mutations in humans was initially believed to be associated with impaired hepatobiliary Mn elimination (35). However, liver-specific Zip14 KO mice do not show systemic Mn accumulation or motor deficits (37), suggesting that Mn is eliminated by an alternative route. In enterocytes, ZIP14 has a basolateral orientation (BL) (3, 14). We have previously shown that when administered subcutaneously, ^54^Mn levels were lower in intestinal tissue and the intestinal contents of whole-body *Zip14* KO (WB-KO) mice compared with WT mice, suggesting that systemic Mn elimination is less effective in the absence of ZIP14 (3). Thus, our working hypothesis centers on BL-localized ZIP14 in enterocytes providing a route for elimination of Mn. Using comparisons between whole-body and intestine specific *Zip14* KO mice, we demonstrate here that deletion of intestinal *Zip14* is sufficient to cause systemic Mn overload and that intestinal ZIP14 is a significant contributor to Mn homeostasis.

## Materials and Methods

### Mice

After weaning, the mice were fed ad libitum a commercial chow diet that contained 93 mg Mn/kg (Harlan 7012) and tap water and were maintained on a 12/12 h, light/dark cycle. Mice were used as young adults (8-16 weeks of age). Both sexes were included in all experiments. Euthanasia was through exsanguination by cardiac puncture under isoflurane anesthesia. Protocols were approved by the both Cornell University and University of Florida Institutional Animal Care and Use Committees. *Whole-body Zip14 KO mice*: The design and validation of the conventional *Zip14* KO (*Zip14*^*-/-*^) (WB-KO) murine strain has been described previously (2). The breeding colony has been maintained with backcrosses of the C57BL/6;129S5 background. The mice used in these experiments were of the F12 generation or later. *Intestine-specific Zip14 KO mice*: Floxed Zip14 mice on the 129Sv background were generated using targeting of introns 4 and 8. Founder *Zip14*^*flox/+*^ mice were bred to obtain *Zip14*^*flox/flox*^ (*Zip14*^*fl/fl*^) mice at the University of Florida. Following genotype confirmation, *Zip14*^*fl/fl*^ mice were crossed with B6.Cg-Tg(Vil1-cre)997Gum/J (Jackson Lab stock # 004586) mice to create an intestine-specific *Zip14* KO (I-KO) mouse model with which to evaluate the role of intestinal ZIP14 on Mn absorption and excretion.

### Treatments

Following four h of morning fasting, mice were administered ^54^Mn either via gavage (5 µCi) or subcutaneous injections (3 µCi). Four h later, the mice were euthanized via cardiac puncture to collect blood and tissues. The entire excised small intestine was perfused with a metal chelating buffer (10 mM EDTA, 10 mM HEPES and 0.9% NaCl2). The ^54^ Mn content of tissues was measured by gamma-ray solid scintillation spectrometry and normalized by tissue weight. *Manganese exposure*: Either control or Mn-supplemented (2mg Mn/L as MnCl_2_) water was provided to the mice for one month. We chose to supplement Mn via the drinking water to maintain a relatively constant consumption of the diet. *Genotyping*. Genomic DNA was extracted from mouse tail biopsies using extraction buffer (25mM NaOH / 0.2 mM EDTA) and incubation at 98ºC for 45 min (33). The genotyping protocol for whole body *Zip14* KO mice has been presented previously (2). Following primer sets were used for genotyping *Zip14*^*fl/fl*^ and Villin-Cre+ mice: Flox-Forward 5’–AGT GGC CAT GGT AGT TCC TG-3’, Flox-Reverse 5 –CCT GGT GCC TGC ATA TTC TC-3’ and Cre-Reverse 5’-CAT GTC CAT CAG GTT CTT GC-3’, Cre- Forward 5’-TTC TCC TCT AGG CTC GTC CA-3’, Internal Positive Control Forward 5’-CTA GGC CAC AGA ATT GAA AGA TCT-3’, Internal Positive Control Reverse 5’-GTA GGT GGA AAT TCT AGC ATC ATC C-3’, respectively.

### Metal Assays

To measure Mn concentrations, weighed aliquots of tissue were digested at 95°C for at least 3 h in HNO_3_. Whole blood (500µL) was dried and then digested for 24 h in HNO_3_. The entire excised small intestine was perfused with a metal chelating buffer (10 mM EDTA, 10 mM HEPES and 0.9% NaCl2) (equal volume/intestine), then digested for 24 h in HNO_3_. Digested samples were diluted in Milli-Q water. Mn was measured by Microwave Plasma-Atomic Emission Spectrometry (MP-AES) using 403.076 nm for emission detection. Normalization was to tissue or body weight (intestinal contents).

### ROS assay

Whole brain tissues were homogenized in lysis buffer (250 mM Sucrose 20 mM HEPES-NaOH, PH: 7.5, 10 mM KCl, 1.5 mM MgCl_2_, 1 mM EDTA, 1 mM EGTA) that supplemented with protease inhibitors. Following centrifugation at 800 x g for 10 min, supernatants were incubated with 2,7-Dichlorodihydrofluorescein diacetate (Cayman 4091-99-0) at 37 ºC for an h. The fluorescent signal was measured at 502/523 nm.

### Western blot

Tissue was homogenized in lysis buffer with protease and phosphatase inhibitors (Thermo Scientific and AGScientific) added along with the PMSF (Sigma-Aldrich) using the Bullet Blender (Next Advances). Solubilized proteins were separated by 10% SDS-PAGE. Visualization was by chemiluminescence (SuperSignal, Thermo Fisher) and digital imaging (Protein Simple). Rabbit anti-mouse ZIP14 antibody was custom made by Genscript. Antibodies for GAPDH and actin were obtained from Cell Signalling.

### Quantitative PCR analyses

Tissues were homogenized in TRIzol reagent using the Bullet Blender homogenizer and RNA was isolated. Total RNA was used to measure relative mRNA abundance by quantitative PCR (qPCR). Primer/probes for iNOS, tnfα, il6, and gapdh were purchased from Applied Biosystems.

### Motor Function

Motor functions were tested in the light-phase. All tests were performed as in previous studies (3). *Beam Traversal*. The beam was constructed of four segments of 0.25 m in length. Each segment was of thinner widths 3.5 cm, 2.5 cm, 1.5 cm, and 0.5 cm; The widest segment acted as a loading platform for the mice, and the narrowest end was adjacent to the home cage. After two days of training, on the test day, mice were timed over three trials to traverse from the loading platform and to the home cage. *Pole Descent*. A pole (0.5 m long, 1 cm in diameter) was placed in the home cage. After two days of training, on the test day, the time to descend from the top of the pole to the cage floor was measured. *Fecal output*. Mice were transferred from their home cage to a 12×20 cm translucent cylinder, and fecal pellets produced over 5 min were counted. The data were generated by using averages of three days of trials. *Inverted grid*. The mice were placed in the center of a 26 by 38 cm screen with 1-cm^2^-wide mesh. The screen was inverted for 60 s, and the mice were timed until they released their grip or held on.

### Magnetic Resonance Imaging

The MRI scans were collected in a 4.7T/33 cm horizontal bore magnet (Magnex Scientific) at the Advanced Magnetic Resonance Imaging and Spectroscopy facility in the McKnight Brain Institute of the University of Florida. The MR scanner consisted of a 11.5 cm diameter gradient insert (Resonance Research, Billerica, MA, USA) controlled by a VnmrJ 3.1 software console (Agilent, Palo Alto, CA, USA). A quadrature transmit/receive radiofrequency (RF) coil tuned to 200.6 MHz ^1^H resonance was used for B1 field excitation and RF signal detection (airmri; LLC, Holden, MA). On the scanning day, mice were induced using 3-4% isoflurane delivered in medical grade air (70% nitrogen, 30% oxygen; air flow rate 1.5 mL/min). The anesthesia was maintained at 1.0-1.5% isoflurane during MRI scanning. Core body temperature and spontaneous respiratory rates were continuously recorded during MRI scanning (SA Instruments, Stony Brook, NY). Mice were maintained at normal body temperature levels (37– 38 °C) using a warm water recirculation system. The MRI included a multiple repetition time sequence to calculate parametric T_1_ maps for each group using a fast spin echo sequence with a total of four TR’s (0.5, 1.08, 2.33, 5.04 seconds), and TE = 6.02 ms with the following geometric parameters: 16 x 16 mm^2^ in plane, 14 slices at 0.8 mm thickness per slice, data matrix = 128 x 128 (125 μm in-plane resolution).

### MRI Post-Processing and analysis

Whole brain masks were obtained via automated segmentation with 3-dimensional pulse-coupled neural networks (PCNN3D) using high-resolution anatomical scans to remove non-brain voxels. All cropped data were used to create templates for each cohort using Advanced Normalization Tools (ANTs; http://stnava.github.io/ANTs/). The templates were then registered to an atlas of the mice brain using the FMRIB Software Library linear registration program flirt. The atlas was then transformed back to each individual data set with the registration matrices from ANTS. To generate parametric T1 maps, multi-TR images were fit to the equation S_TR_=S_0_(1-e^−TR/T1^) using non-linear regression in ImageJ. From T1 maps, the T1 relaxation rate (R1 in ms^−1^) is calculated and exported from regions of interest.

### Statistics

For all experiments, both sexes of mice were included. Data are presented as means <u>± SD</u>. Significance was assessed by Student’s t-test for single comparisons and by ANOVA/Tukey’s test for multiple comparisons. Statistical significance was set at P<0.05. Analyses were performed using GraphPad Prism.

## Results

### Deletion of intestinal *Zip14* causes impaired serosal to mucosal ^54^ Mn transport

A representative PCR amplification for genotyping was shown in Fig. 1A. Intestine-specific ablation of *Zip14* was confirmed by western blotting of ZIP14 protein. Tissues from both VC and fl/fl mice were included as controls. ZIP14 was deleted only in the intestine of the I-KO mice (Fig. 1B). Western blotting shows expression of ZIP14 between I-KO and WB-KO mice (Fig. 1C). Absence of ZIP14 from only the intestine of the I-KO mice confirms tissue specificity. To determine the role of intestinal ZIP14 in Mn clearance, we conducted ^54^Mn uptake studies using I-KO mice to assertain the direction of Mn transport mediated by ZIP14. ^54^Mn was administered via two routes, and radioactivity in the intestine was measured. There was no difference in the amount of radioactivity in the intestine of I-KO, VC, and fl/fl mice when ^54^Mn was administered via oral gavage (Fig. 1D), indicating that intestinal ^54^Mn absorption was not affected by ZIP14 deficiency. However, significantly (p<0.001) less radioactivity was detected in the intestine of *Zip14* I-KO mice when ^54^Mn was administered via sc injection, indicating that the intestine-specific deletion of *Zip14* decreased serosal to mucosal transcellular Mn transport. In the following experiments, fl/fl mice were used as controls since we have confirmed that there was no difference in ^54^Mn uptake between fl/fl and VC mice (Fig. 1D).

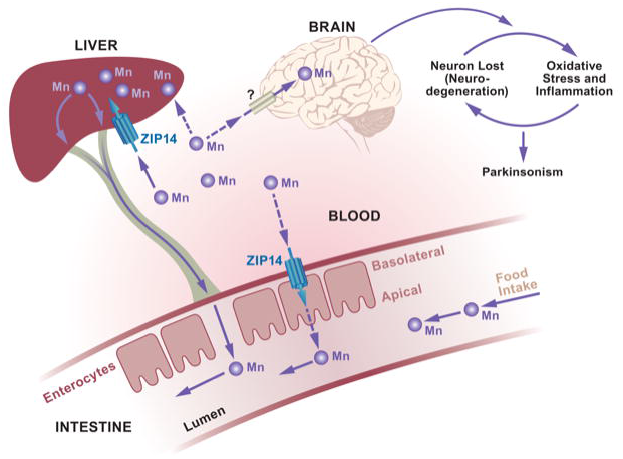

**Figure 1.**
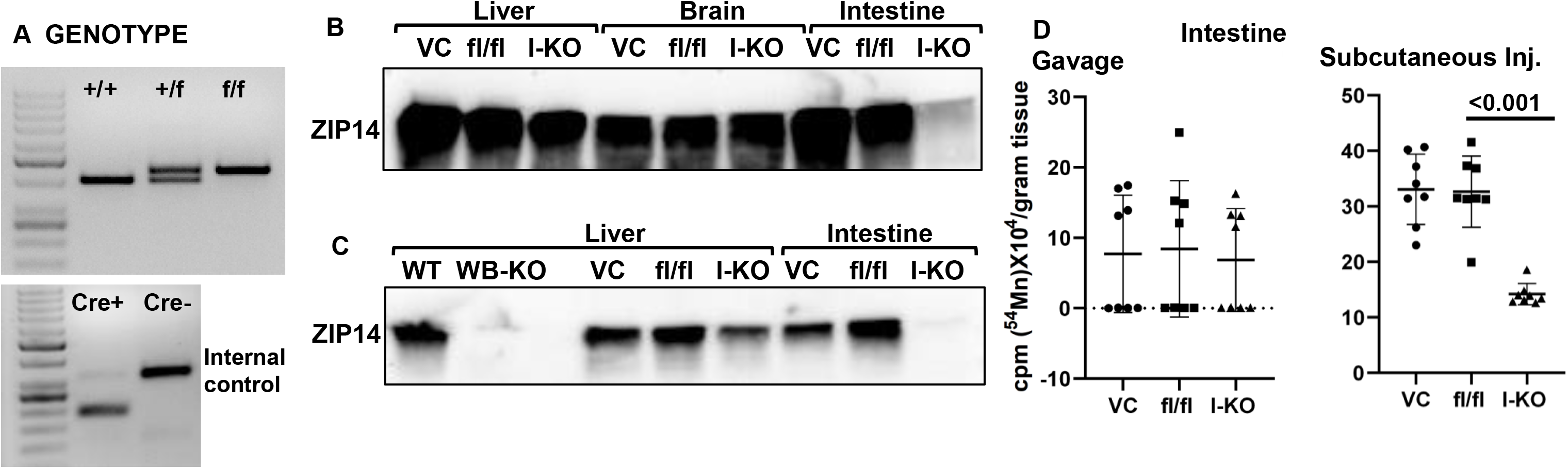
^54^Mn elimination is impaired in intestine-specific *Zip14* KO mice. A) Representative genotype B) Representative western analyses are showing ZIP14 protein levels. B) ZIP14 protein levels in liver, brain, and intestine of the villin-cre (VC), floxed *Zip14* (fl/fl) and intestine-specific conditional *Zip14* KO (I-KO) mice C) Comparison of ZIP14 protein levels between whole-body and intestine-specific *Zip14* KO. D) Mice were administered ^54^Mn via either gavage or subcutenous injection. Four h later, intestinal 54Mn absorption and elimination were measured in floxed *Zip14* (fl/fl) and intestine-specific *Zip14* KO (I-KO) mice. Values are means +/− SD; n = 8 (both female and male mice were included). Student’s t test for fl/fl vs. I-KO comparison.

### Deletion of intestinal *Zip14* leads to systemic and brain Mn accumulation

To test whether impaired intestinal ZIP14-mediated Mn excretion affects systemic Mn metabolism, we compared systemic ^54^Mn distribution between WB-KO and I-KO mice. Our results revealed that when it was given via s.c. injection, the amount of ^54^Mn was significantly less in the intestines of both WB- and I-KO mice (Fig. 2A). Furthermore, the amount of ^54^Mn was significantly greater in the blood and brain of the WB and I-KO mice when compared to their controls (Fig. 2B). The magnitude of increase in ^54^Mn was less in I-KO mice compared to WB-KO mice.

**Figure 2.**
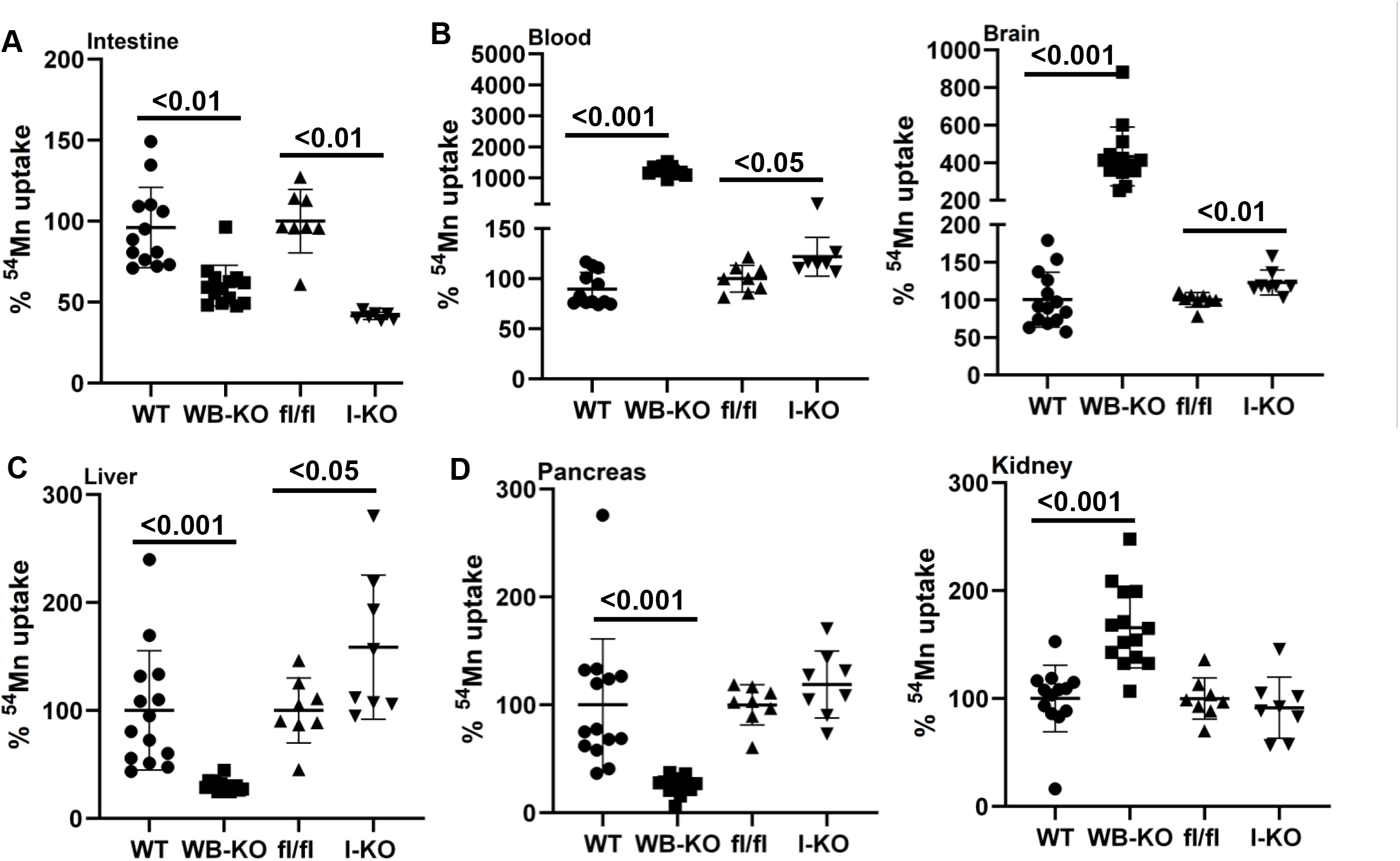
Comparison of percent ^54^Mn uptake between whole-body and intestine-specific *Zip14* KO. Mice were administered ^54^Mn via subcutenous injection, and 4 h later radioactivity was measured in intestine (A), blood and brain (B), liver (C), and pancreas and kidney (D). Values are reported as the mean+/-SD. N=7–14 (both female and male mice were included). Student’s t test for WT vs WB-KO and fl/fl vs I-KO comparison.

The key route of Mn elimination is through the hepatobiliary system. To evaluate the hepatobiliary clearance of Mn, we measured liver ^54^Mn uptake. Of note ^54^Mn uptake was impaired in WB-KO and enhanced in I-KO, suggesting that a lack of intestinal ZIP14 was compensated for in the I-KO mice through increased Mn elimination via the hepatobiliary system (Fig. 2C). Furthermore, ^54^Mn uptake was not different between kidney and pancreas of fl/fl and I-KO mice, while ^54^Mn uptake was higher and lower in pancreas and kidney of WB-KO, respectively suggesting that ZIP14 might be involved with ^54^Mn uptake in the kidney (Fig. 2D).

Next, to evaluate the influence of intestinal ZIP14 on brain Mn, we measured Mn accumulation using MP-AES. The Mn concentration was greater in the blood and brain of both the WB-KO and I-KO mice compared to their controls (Fig. 3A). As expected, relative accumulation was less in the I-KO mice. Furthermore, we used MRI analysis to evaluate brain Mn accumulation between WB-KO and I-KO mice. Signal intensity was greater in the brains of I-KO when compared to fl/fl. Similar to ^54^Mn uptake and MP-AES data, the differences in MRI signal intensity between the brains of the fl/fl and I-KO were less than that of between the WB-KO and WTmice (Fig. 3B). Nevertheless, the images from the I-KO mice clearly show punctate regions of increased intensity consistent with Mn accumulation. Of note, the MRI images reflected endogenous Mn in brains of both genotypes obtained without any enhancement and were from young adult mice maintained under barrier conditions with normal husbandry. These data collectively suggest that deletion of intestinal *Zip14* was sufficient to cause systemic Mn overload and that intestinal ZIP14 is a significant contributor to Mn homeostasis.

**Figure 3.**
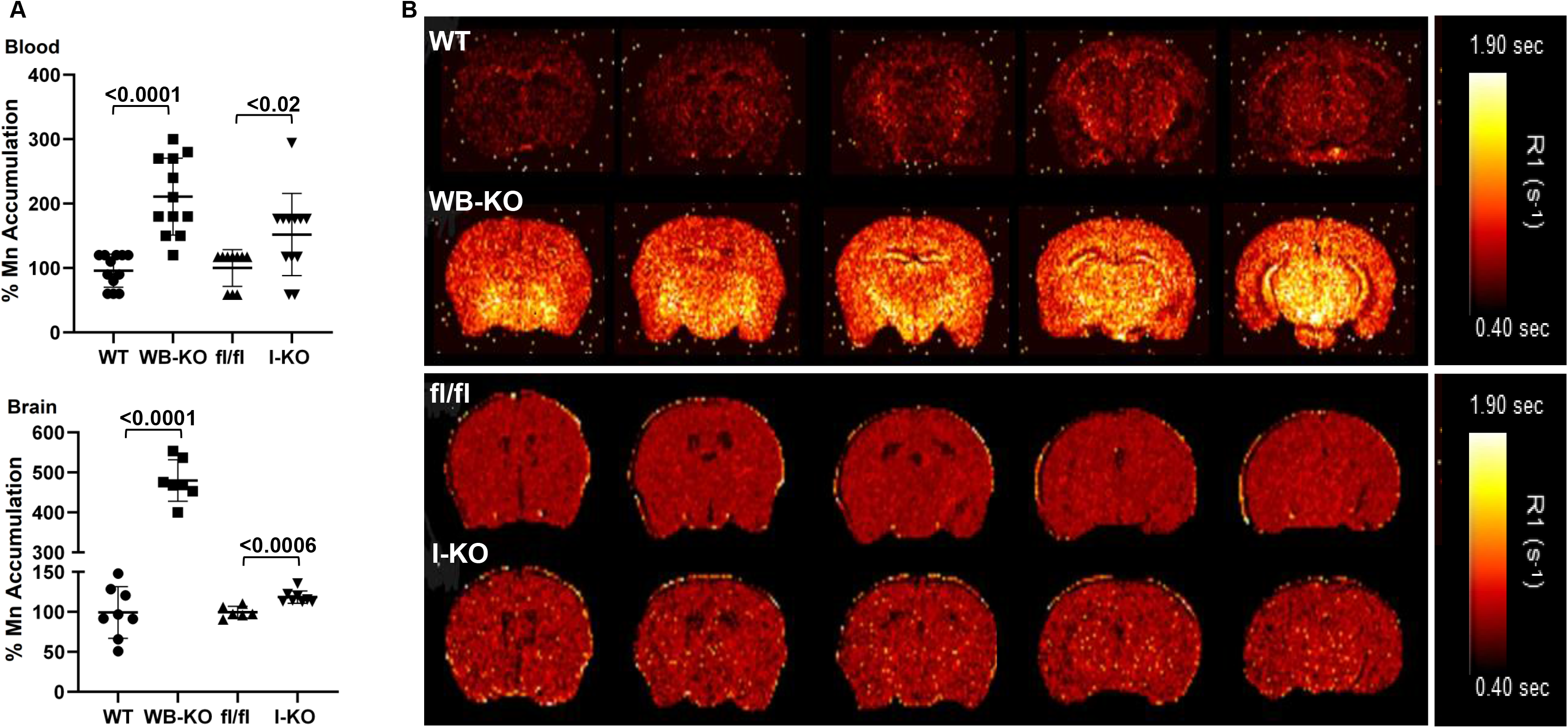
Deletion of intestinal *Zip14* leads to manganism in mice. A) Mn concentrations in blood and brain were measured by Microwave Plasma-Atomic Emission Spectrometer (MP-AES). B) Representative MR images with a quantitative map of R1 relaxation rates (inverse of T1 values).Values are reported as the mean+/-SD. N=7-14 (both female and male mice were included). Student’s t test for WT vs WB-KO and fl/fl vs I-KO comparison.

### Neuroinflammation and motor dysfunction in intestine-specific *Zip14* KO is amplified with high Mn exposure

Deletion of intestinal *Zip14* caused systemic and brain Mn overload; however, the magnitude of increases was less in I-KO compared to WB-KO. Therefore, we have conducted Mn supplementation studies with I-KO mice to increase exposure. Deletion of intestinal ZIP14 was confirmed by western blot (Fig. 4A). Following Mn-exposure, increased intestinal Mn concentrations were found in the fl/fl mice, but were not observed in the I-KO mice (Fig. 4A). Moreover, the Mn concentration of luminal and fecal content of Mn-supplemented I-KO was significantly less than Mn-supplemented fl/fl mice (Fig. 4B). Mn concentrations were greater in the liver of I-KO mice compared fl/fl mice in the control group demonstrating greater hepatic Mn clearance occurs when hepatic ZIP14 is functional and intestinal ZIP14 is not (Fig. 4C).

**Figure 4.**
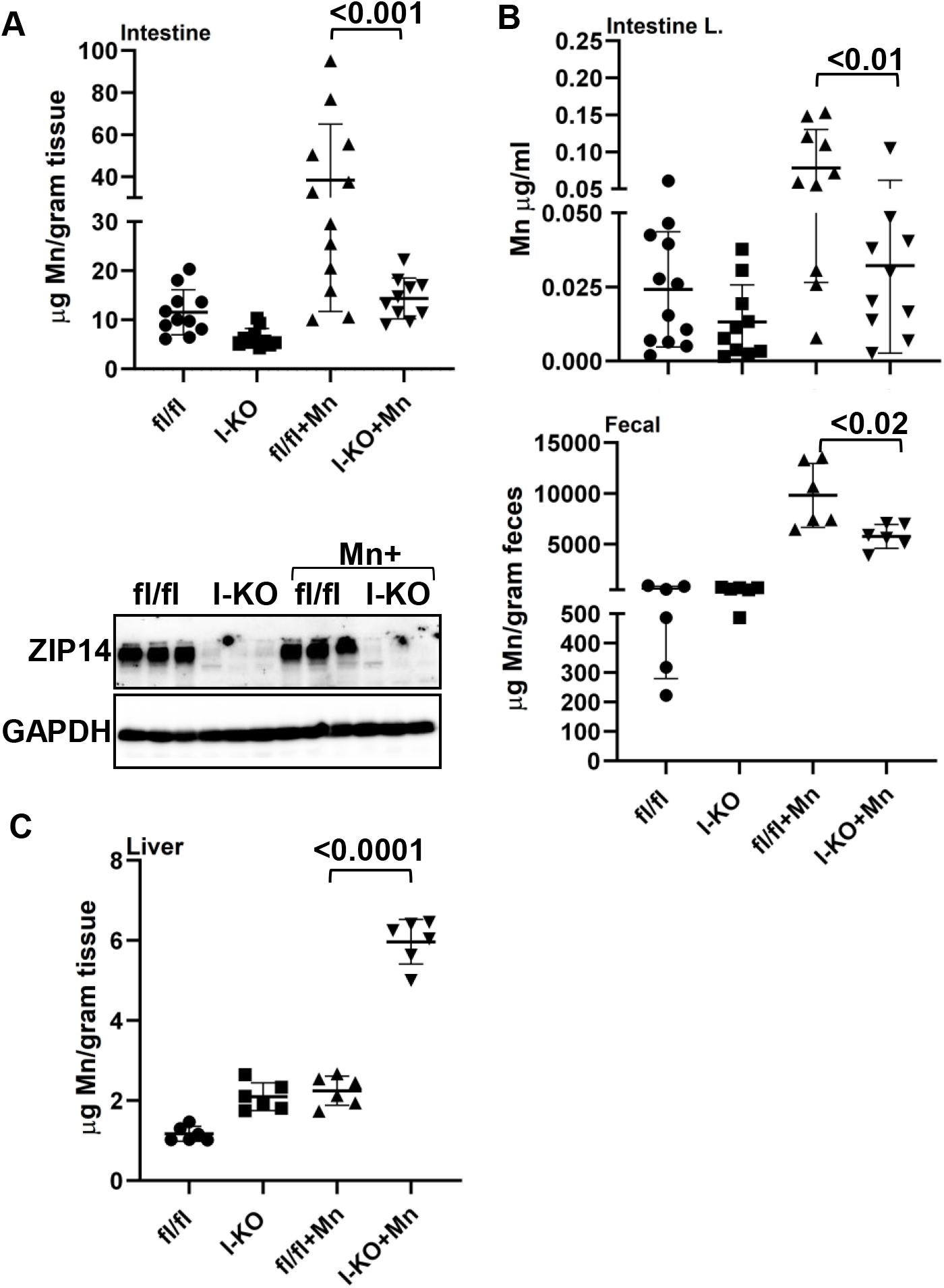
Comparison of Mn concentration between control and Mn-supplemented intestine-specific *Zip14* KO. A) Representative western blot images showing the absence of ZIP14 in I-KO intestines. Mn concentrations in the intestine (A), intestine luminal and fecal content (B) and liver (C) were measured by MP-AES. Values are means+/− SD. n=6-12 (both female and male mice were included). Significance was assessed by One-way ANOVA.

Mn is neurotoxic; thus, we next measured the blood and brain Mn concentrations. Mn concentrations were greater in blood and brain of I-KO mice compared fl/fl mice in the control group (Fig. 5A, B). Mn concentrations in Mn-supplemented fl/fl mice were comparable to the I-KO unsupplemented mice. Mn-supplementation further enhanced the Mn accumulation in blood and brain of I-KO mice compared to Mn-supplemented fl/fl mice, however. Importantly there was no change between groups in brain Zn and Fe concentrations (Fig. 5C). Importantly, the Mn-supplemented I-KO group had the greatest increase in Mn concentrations, further confirming impaired Mn excretion in I-KO mice occurs via the intestine.

**Figure 5.**
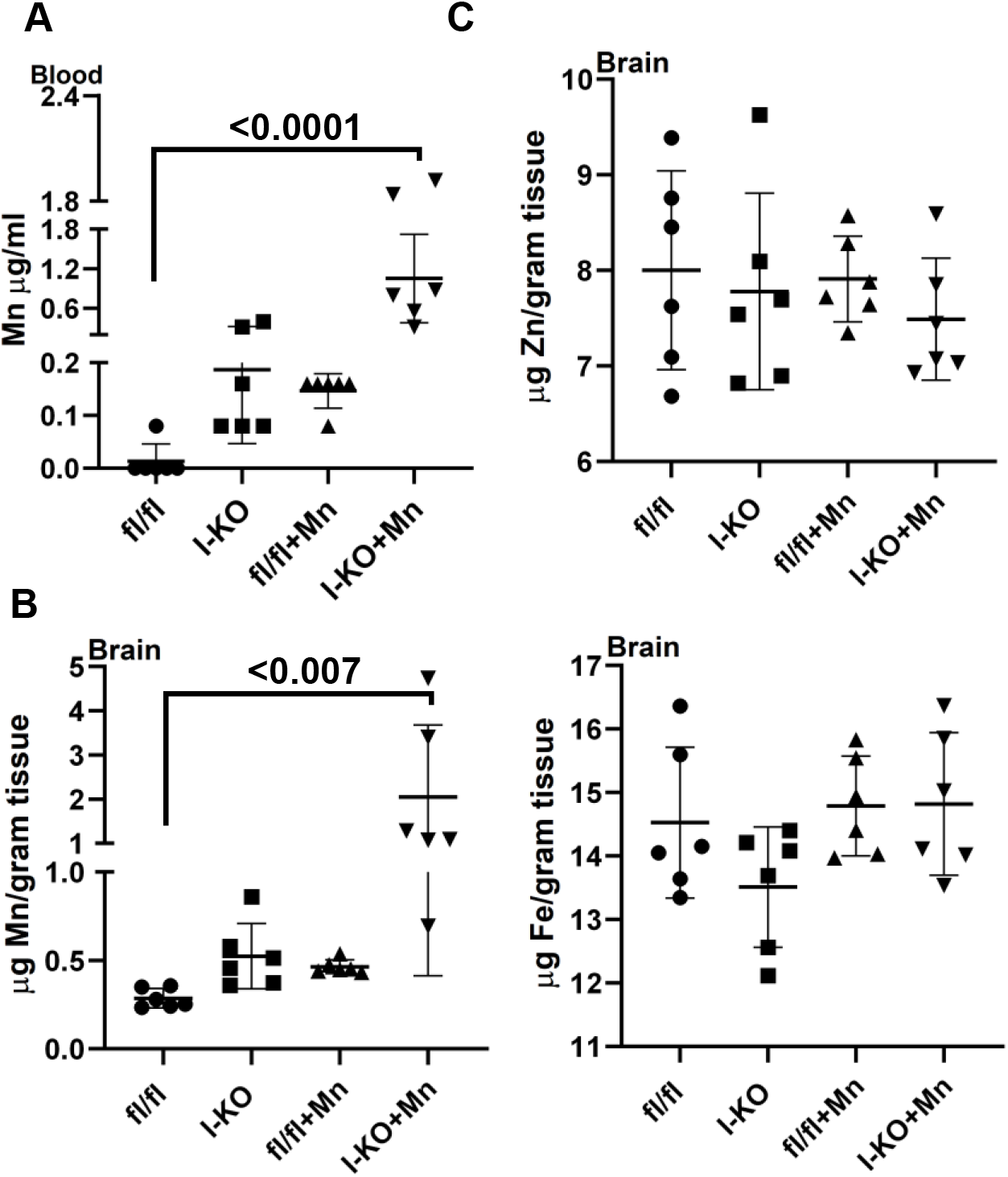
Comparison of metal concentrations between control and Mn-supplemented intestine-specific *Zip14* KO. Mn concentrations in blood (A) and brain (B), and Zn and Fe concentrations in the brain (C) were measured by MP-AES. Values are means+/− SD. n=6 (both female and male mice were included). Significance was assessed by One-way ANOVA.

To test the physiological consequences of Mn overload, we assessed motor functions using the inverted grip, balance beam traversal, pole descent test, and fecal output methods. The Mn-supplemented I-KO mice required significantly more time to cross a balance beam compared to fl/fl control mice (Fig. 6A) while there was no difference in the time to descend a pole (data not shown). Furthermore, the total output of fecal pellets was significantly lower in Mn-supplemented I-KO mice (Fig. 6A). We found no difference in body weight and limb strength (inverted grid assay) (data not shown). These data collectively showed that at the end of one month of chronic Mn exposure, I-KO mice developed initial indices of motor dysfunction; however, the whole spectrum of manganism had not been obtained.

**Figure 6.**
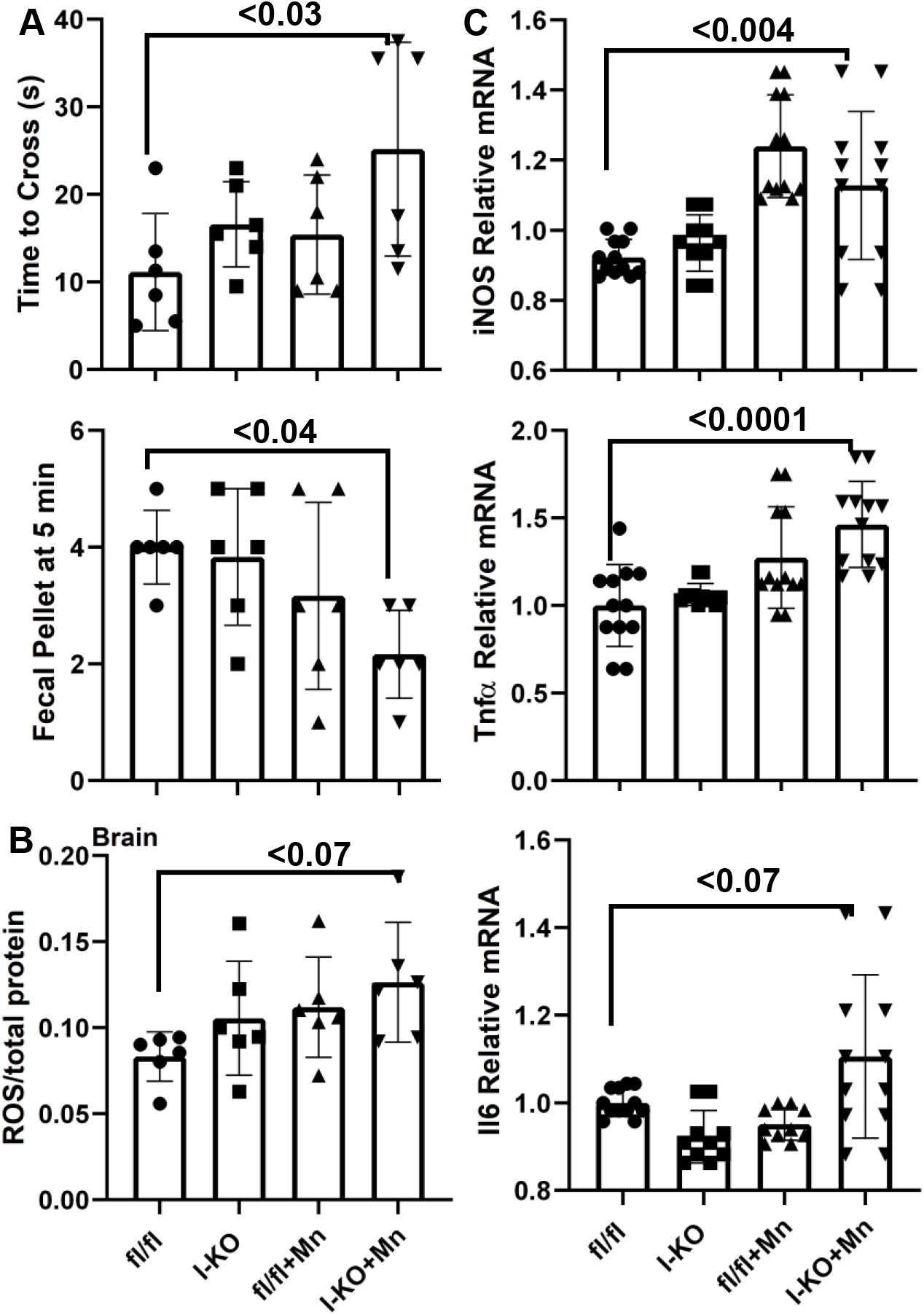
Comparison of motor functions between control and Mn-supplemented intestine-specific *Zip14* KO. A)Time to cross the balance beam and fecal output were counted. ROS (B) and cytokine expressions (C) were measured. Values are means+/− SD. n=6 (both female and male mice were included). Significance was assessed by One-way ANOVA.

To investigate the biochemical and physiological underpinnings of Mn-induced motor dysfunction, we measured reactive oxygen species (ROS) and cytokine expressions in whole-brain lysates. Our results revealed an increased amount of ROS in the I-KO and Mn-supplemented fl/fl mice compared to fl/fl mice (Fig. 6B). The highest increase was found in Mn-supplemented I-KO mice; the difference was not statistically significant (p=0.07). Similarly, the highest increase for il6 mRNA was found in Mn-supplemented I-KO mice; the difference was not statistically significant (p=0.07) (Fig. 6C). However, the expression of iNOS and tnfα mRNAs was significantly increased in Mn-supplemented I-KO mice when compared to control fl/fl mice (Fig. 6C) suggesting that Mn-induced increase in ROS and cytokines could be an underlying cause for the initiation of parkinsonism. and il6

## Discussion

In the present study, using whole-body and intestine-specific *Zip14* KO mice, we demonstrate that intestinal ZIP14 is essential for Mn homeostasis. Mice with a deletion of intestinal *Zip14* displayed a phenotype that included impaired serosal to mucosal Mn transport resulting in increased liver, blood, and brain Mn accumulation. Mn supplementation greatly enhanced the brain Mn overload in I-KO mice with early parkinsonism symptoms.

Manganese metabolism is homeostatically maintained through limited absorption by the small intestine with elimination occurring via hepatobiliary, pancreatic, and urinary (minimally) excretions (7). The plethora of transporters with proposed roles in manganese metabolism has been documented (15). The list includes divalent metal transporter 1 (DMT1), transferrin receptor (TfR), calcium channels, Park9/ATP13A2, NCX, SPCA1, and ferroportin (FPN). However, the precise molecular mechanisms that underlie Mn homeostasis are poorly understood. ZIP14/SLC39A14 is a ZIP family transmembrane protein that regulates intracellular levels of zinc (Zn), Mn, and iron (Fe). Although occupational and environmental Mn exposures are known sources of Mn toxicity, genetic mutations in metal transporters, including ZIP14, also cause Mn-induced parkinsonism. Mutations in the *SLC39A14* gene encoding the ZIP14 transporter were found in patients with childhood-onset parkinsonism and dystonia with systemic and brain Mn accumulation (35). Deletion of Zip14 expression in zebrafish recapitulated the human brain Mn accumulation phenotype of ZIP14 mutation carriers (35). However, Mn concentration in the abdominal viscera of WT and mutant zebrafish was not different. Using whole-body *Zip14* KO mice we have previously shown that WB-*Zip14* KO mice displayed phenotypes similar to those of human carriers of *ZIP14* mutations, including spontaneous systemic and brain Mn overload with motor dysfunction and impaired hepatic Mn uptake (3). Our findings with *Zip14* KO mice were confirmed by two independent research groups (16, 37) collectively supporting the use of *Zip14* KO mouse models as a relevant model system with which to further investigate the organ/tissue-specific role of ZIP14-mediated Mn transport in Mn homeostasis.

The early studies with ^54^Mn revealed that hepatobiliary system was a major route for Mn elimination from the body (5). However, using liver-specific *Zip14* KO mice, Xin et al. showed that there was no Mn accumulation in the brain or other tissues of L-KO mice, which have considerably lower hepatic Mn levels. This finding supports our hypothesis that there are other highly efficient Mn elimination routes in the body. We previously showed that ZIP14 is localized at the basolateral site of the enterocytes (3, 14). Furthermore, intestinal Mn elimination was impaired in WB *Zip14* KO mice. Therefore we generated I-KO mice to explore the specific role of intestinal *Zip14* in Mn elimination/detoxification. We found that Mn transport in the serosal to mucosal direction was impaired and importantly Mn accumulated in the blood and and brain at steady state. This novel finding suggests that deletion of intestinal *Zip14* is sufficient to cause systemic Mn overload and that intestinal ZIP14 is a significant contributor to Mn homeostasis.

The known Mn elimination routes are hepatobiliary (major), pancreatic, and urinary (minimally) excretion (7). Therefore, we compared ^54^Mn transport in these tissues of WB- and I-KO mice. Pancreatic ^54^Mn uptake was impaired, and ^54^Mn uptake by kidney was enhanced in WB-*Zip14* KO. In contrast, we have not observed any changes in pancreas and kidney of I-KO mice compared to fl/fl control mice. However, there was a significant increase in the liver ^54^Mn uptake and Mn accumulation. This suggests there is an activation of a hepatic compensation mechanism. These novel differences in tissue ^54^Mn uptake and Mn accumulation in I-KO mice are providing new aspects for the paths of systemic Mn detoxification.

Impaired manganese (Mn) homeostasis results in excess Mn and neurotoxicity; however, the underlying molecular mechanisms have not been clearly defined. It has been suggested that oxidative stress and associated neuroinflammation lead to neurodegeneration and parkinsonism (motor dysfunction) (26, 27, 30, 32). Our results revealed that ROS and cytokine expressions were highly upregulated in the brain of Mn-supplemeted I-KO mice. Notably, only Mn (but not zinc or iron) levels in the brains of I-KO and Mn-supplemented mice were greater proving evidence for the specific effect of Mn on neuroinflammation. The main site of Mn-induced inflammation in the brain remains to be elucidated.

In conclusion, we show here the role of intestinal ZIP14 on Mn elimination and homeostasis at the organism level. Mn elimination by intestinal ZIP14 is essential to maintain Mn homesatsis since the hepatic compensation mechanism is not sufficient to prevent Mn overload and subsequent neurotoxicity.

## Acknowlegments

This project was supported by the National Institute of Diabetes and Digestive and Kidney Diseases 5R01-DK 094244 to RJC and Cornell University Division of Nutritional Sciences funds to TBA. The authors acknowledge the support from the National High Magnetic Field Laboratory’s Advanced Magnetic Resonance Imaging & Spectroscopy (AMRIS) Facility (National Science Foundation Cooperative Agreement No. DMR-1157490 and the State of Florida). Graphical abstract was prepared by Allison A. Stevens, AASarts.com.

## Author contributions

T.B.A., and R.J.C. designed research; T.B.A., T.L.T., C.H.R., and M.P. performed research; T.B.A., M.F., and R.J.C. analyzed data; T.B.A. and R.J.C. wrote the paper.

